# Variable expression of fluorescent protein expression in hematopoietic cells purified from ubiquitous reporter mice

**DOI:** 10.64898/2026.11.24.765905

**Authors:** Quan Gu, Daniel Lucas

## Abstract

Adoptive transfer experiments (e.g., hematopoietic stem cell transplantation) are indispensable for hematology and immunology research. These assays require unambiguous detection of the transferred cells, and their offspring, to track their fates. Mice in which a fluorescent reporter protein is expressed ubiquitously in all cells are especially helpful for these fate mapping experiments as the donor cells can be easily detected using flow cytometry and confocal microscopy. We report that two commonly used reporter strains, *Ubc-gfp* and *ROSA-mTmG*, display variable expression during hematopoietic differentiation. GFP in *Ubc-gfp* mice labels only a fraction of megakaryocytes while TdTomato expression in *ROSA-mTmG* mice is progressively lost in red blood cells. We also found uneven expression of either reporter at various stages of B cell, myeloid cell or dendritic cell differentiation. Our results thus enable selection of the appropriate fluorescent reporter for identification of the hematopoietic cells of interest in adoptive transfer experiments.

## Introduction

The hematology field has widely used the CD45.1/CD45.2 congenic system, in which donor and host cells can be distinguished based on the expression of two isoforms of the CD45 protein, to track hematopoietic stem cell engraftment (1-6). The main limitation of this model is that CD45 is not expressed in erythroid cells or platelets, thus precluding evaluation of donor cell contribution to these lineages. To overcome this limitation the field has used mouse models like *Ubc-gfp* mice in which GFP protein is reported to be expressed ubiquitously (7) and allow quantification of donor cell engraftment in all major hematopoietic lineages in the blood, including platelets and red blood cells, as well as other tissues (8-11). Here we report unexpected loss of GFP expression in megakaryocytes from *Ubc-gfp* mice. Instead, the contribution of donor cells to megakaryocytes can be efficiently tracked by detecting mTdTomato expression in *Rosa-mTmG* mice, another model of ubiquitous fluorescent labeling (12). Surprisingly, we found variable reporter fluorescent protein expression at different stages of hematopoiesis in both *Ubc-gfp* and *Rosa-mTmG* mice. We provide a blueprint for optimal fluorescent reporter model selection to track specific cell populations in the bone marrow in adoptive transfer experiments.

## Materials and Methods

### Mice

Mice were housed under a 12-hour light and 12-hour dark cycle in specific pathogen-free conditions with ambient temperature (22 °C) and humidity (40–60%) in a dedicated vivarium at Cincinnati Children’s Hospital Medical Center. All experimental procedures were approved by the Institutional Animal Care and Use Committee (IACUC) of Cincinnati Children’s Hospital Medical Center. We used the following mouse strains purchased from the Jackson laboratory: B6.SJL-*Ptprcapepcb*/BoyJ (CD45.1^+^), C57BL/6-Tg (UBC-GFP)30Scha/J *(Ubc-gfp*) (7) and B6.129(Cg)-*Gt (ROSA)26Sortm4(ACTB-tdTomato, -EGFP)Luo*/J (*mT*/mG) (12).

### Bone marrow and blood collection

Mice were euthanized by isoflurane inhalation followed by cervical dislocation. Bone marrow cells were harvested by flushing bones into with 1 mL of pre-chilled PEB buffer (2 mM EDTA and 0.5% bovine serum albumin in phosphate-buffered saline (PBS)). Blood was collected from the retro-orbital venous sinus into EDTA-coated tubes. For analysis of non-erythroid cells in the blood, samples were treated with red blood cell (RBC) lysis buffer (150 mM NH_4_Cl, 10 mM NaCO_3_ and 0.1 mM EDTA), centrifuged, resuspended in pre-chilled PEB buffer, and used for subsequent assays.

### *Flow* cytometry

Cells were stained with the indicated antibodies for 30 minutes in dark, washed thrice with pre-chilled PEB buffer, and analyzed in a BD LSRFortessa™ Flow Cytometer (BD Bioscience). Dead cells and doublets were excluded based on forward scatter/side scatter (FSC/SSC) profiles and 4′,6-diamidino-2-phenylindole (DAPI) exclusion (Sigma-Aldrich). Antibodies used were: B220 (clone RA3-6B2), CD2 (clone RM2-5), CD3e (clone 145-2C11), CD4 (clone RM4-5), CD5 (clone 53-7.3), CD8 (clone 53-6.7), CD11b (clone M1/70), CD11c (clone N418), CD16/32 (clone 93), CD24 (clone 30-F1), CD41 (clone MWReg30), CD42d (clone 1C2), CD43 (clone S11), CD45 (clone 30-F11), CD48 (clone HM48-1), CD71 (clone RI7217), CD105 (clone MJ7/18), CD115 (clone AFS98), CD127 (clone A7R34), CD135 (clone A2F10), CD150 (clone TC15-12F12.2), CD170 (clone S17007L), CD172a (clone P84), FcεRIα (clone MAR-1), Gr-1 (clone RB6-8C5), IgD (clone 11-26 c.2a), IgM (clone RMM-1), Ly6C (clone HK1.4), Ly6G (clone 1A8), NK 1.1 (clone PK136), Sca-1 (clone D7), Ter119 (clone TER-119), MHCII (clone M5/114.15.2), from BioLegend; CD34 (clone RAM34) and CD117 (clone 2B8), from BioLegend or Thermo Fisher Scientific; CD71 (clone C2) from BD Bioscience. Panel-specific lineage antibody cocktails were applied to gate out unwanted cell populations. HSC/MPP panel: B220, CD3e, CD4, CD5, CD8, Gr-1, CD127 and CD11b; B lymphopoiesis panel: CD2, CD3e, CD5, CD8a, CD11b and Ly6G; other progenitor panel: B220, CD3e, Gr-1 and CD11b; DC-poiesis panel: CD3, Ly6G, Ter119, B220, CD11c and CD11b (for CDP); CD3, B220, NK1.1, Ly6G and Ter119 (for Pre DC); CD3, NK1.1, Ly6G and Ter119 (for cDC and pDC). Data were analyzed with BD FACSDiva™ Software (BD Biosciences) and FlowJo (Tree Star). Gating strategies are shown in Supplementary Figure 1.

### Confocal microscopy and image analyses

Whole-mount sternum immunostaining was performed as described (13). Confocal imaging was performed using Nikon AXR Inverted Confocal Microscope with the following specifications: high power 488 nm, 594 nm and 640 nm. Images were acquired using a LWD Lambda S 40× C water-immersion objective (Nikon, MRD77410) with a resonant scanner and 4× line averaging, 1024 × 1024 pixels resolution. Original images (. nd2 format file) were denoised by an artificial intelligence algorithm (Denoise.AI), stitched to large image using the NIS-Elements software (Nikon, version 5.30.03) and converted to Imaris (.ims) files using Imaris software (Bitplane, version 10.0) for further analyses.

### Statistical analyses

All statistical analyses were performed using Prism 10 (GraphPad Software). Normally distributed data were analyzed using an unpaired two-tailed Student’s *t*-test while Mann-Whitney test was applied to non-normally distributed data.

### Data sharing

For Fig. 1, image files are available at https://doi.org/10.17632/hyhg9h6rwc.1, https://doi.org/10.17632/7svmjr3wnk.1, https://doi.org/10.17632/6db8mxy3zy.1, https://doi.org/10.17632/ts3hfpg76f.1, https://doi.org/10.17632/h6n4sc9kbr.1, https://doi.org/10.17632/9mfzhztb4j.1, https://doi.org/10.17632/6nrsjj55pw.1, https://doi.org/10.17632/gk6hr2b6hw.1, https://doi.org/10.17632/9vw4bfw97v.1, https://doi.org/10.17632/zt736zkpx4.1 and https://doi.org/10.17632/tw5p5kwvzk.1. All flow cytometry and quantification datasets (flow cytometry and imaging) are available at https://doi.org/10.17632/9xmnrds9wr.1.

**Figure 1.**
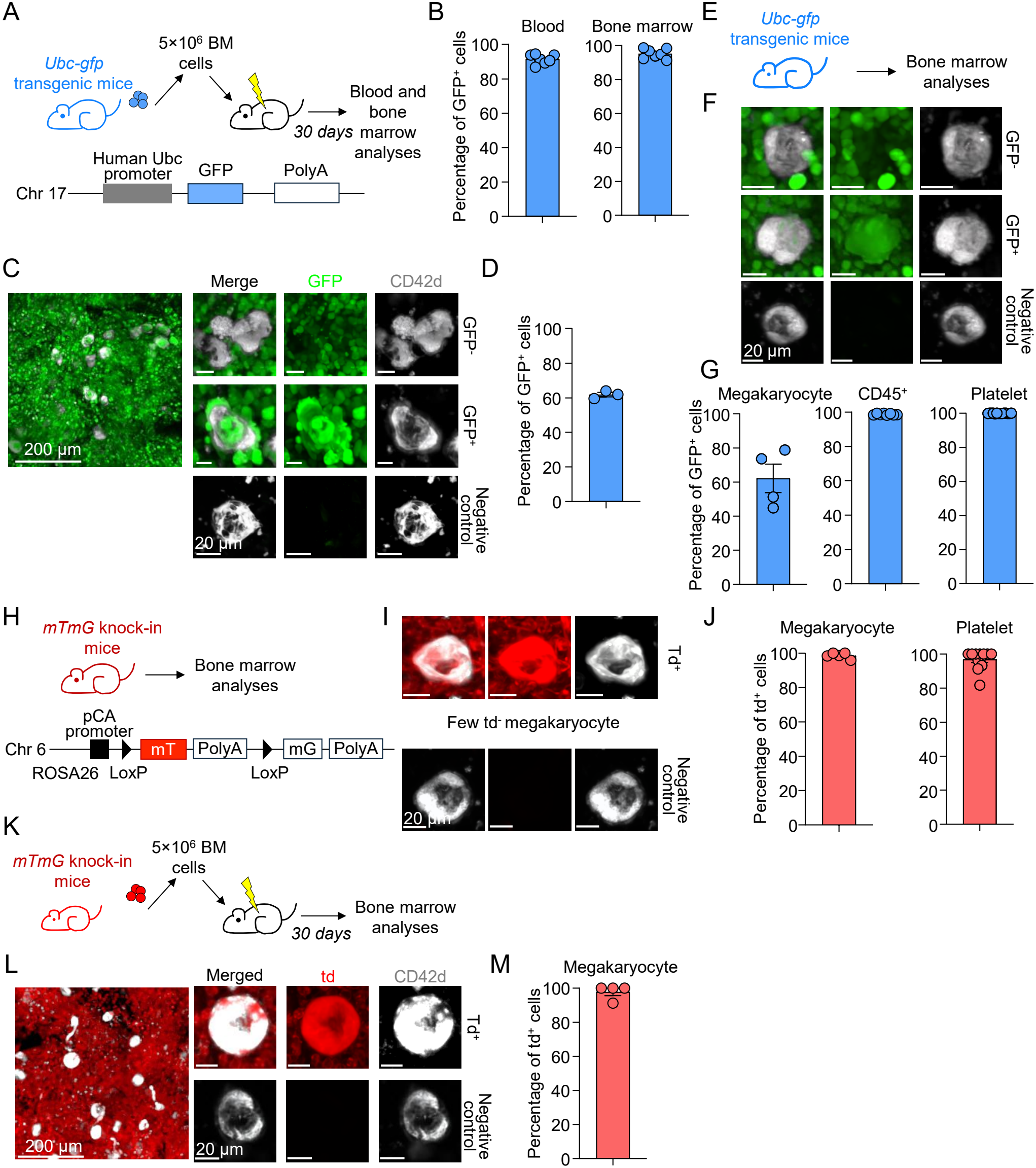
Heterogenous GFP expression in megakaryopoiesis from *Ubc-gfp* mice. (A) Scheme of bone marrow transplantation of GFP^+^ cells from *Ubc-gfp* mice as donor into B6.SJL-*Ptprcapepcb*/BoyJ (CD45.1^+^) congenic recipients. (B) Peripheral blood and bone marrow GFP^+^ cell frequency in single live cells on day 30 after transplantation. (C-D) Representative images (C) and quantification (D) of GFP^+^ and GFP^-^ CD42d^+^ megakaryocytes on day 30 after transplantation. (E-G) Scheme (E), representative images (F), and quantification (G) showing that bone marrow hematopoietic cells and blood platelets in *Ubc-gfp* mice under steady state are uniformly GFP^+^ but only 60% of megakaryocytes are GFP^+^. (H-J) Scheme (H), representative images (I) and quantification (J) of TdTomato expression in megakaryocytes and platelet in *mT/mG* mice under steady state. pCA: CMV enhancer/chicken beta-actin core promoter. (K) Scheme of bone marrow transplantation of TdTomato^+^ cells from *mTmG* mice as donor into CD45.1^+^ congenic recipients. (L-M) Representative images (L) and quantification (M) of megakaryocyte on day 30 after transplantation. In all panels expression of fluorescent proteins was detected by confocal microscopy for megakaryocytes and FACS for all other hematopoietic cells. Each dot represents one mouse in at least two independent experiments.

## Results

Our group has a long-standing interest in understanding the anatomy of hematopoiesis in homeostatic and stress conditions (14). To start dissecting the anatomy of regenerative hematopoiesis we adoptively transferred whole bone marrow cells from *Ubc-gfp* donor mice into lethally irradiated B6.SJL-*Ptprc*^*a*^*pepc*^*b*^/BoyJ (CD45.1^+^) congenic recipients and examined the blood and bone marrow 30 days later (Fig. 1A). As predicted, the recipient mice showed >90% GFP^+^ (donor) cells in the peripheral blood and bone marrow (Fig. 1B). Unexpectedly, only 60% of the megakaryocytes in the bone marrow were GFP^+^ (Fig. 1C, D). The most likely explanations for this observation are that host megakaryocytes (GFP^-^) persist after transplantation or that GFP expression in megakaryocytes is heterogeneous. To distinguish between these possibilities, we examined the bone marrow of eight-week-old *Ubc-gfp* mice in homeostasis (Fig. 1E). We found that only 60% of megakaryocytes in the *Ubc-gfp* bone marrow were GFP^+^ (versus approximately 100% of CD45^+^ cells, Fig. 1F, G). These results were especially surprising because the platelets of *Ubc-gfp* mice were uniformly GFP^+^ (Fig. 1G) suggesting that the subset of GFP^-^ megakaryocytes does not produce platelets or that the platelets produced by these megakaryocytes quickly accumulate GFP after being generated. Together these experiments also demonstrate that *Ubc-gfp* mice should not be used to fate map megakaryocytes in adoptive transfer experiments. To overcome this limitation, we decided to examine *Rosa-mTmG* mice, another reporter model in which a construct containing CMV enhancer/chicken beta-actin core promoter (pCA) and mTdTomato-mGFP cassette was inserted in the ROSA26 locus in chromosome 6 (12). The mTdTomato and polyA are flanked by LoxP sites (Fig. 1H). Cre-mediated recombination results in irreversible loss of TdTomato expression and *de novo* expression of membrane bound GFP. We found homogeneous TdTomato expression in the bone marrow megakaryocytes as well as peripheral blood platelet from *Rosa-mTmG* mice (Fig. 1I, J) demonstrating that this model can be used to efficiently track donor-derived megakaryocytes in adoptive transfer experiments. We then injected total bone marrow cells from *Rosa-mTmG* mice into lethally irradiated CD45.1^+^ congenic recipients and examined the bone marrow on day 30 post transplantation (Fig. 1K). Imaging analyses showed that almost 100% of megakaryocytes are TdTomato^+^ (Fig. 1L, M), indicating that TdTomato from *Rosa-mTmG* mice could maintain long term and stable expression in megakaryocyte.

The results in Figure 1 prompted us to examine whether expression of fluorescent proteins in *Ubc-gfp* or *Rosa-mTmG* mice varied during hematopoietic development. For these experiments we compared GFP or TdTomato expression in eight-week-old *Ubc-gfp* or *Rosa-mTmG* heterozygous mice in bone marrow hematopoietic cells by flow cytometry (Supplementary Fig. 1). Both fluorescent proteins were uniformly expressed at high levels in stem cells and multipotent progenitors, myeloid progenitors, and all steps of neutrophil and monocyte maturation (Supplementary Fig. 2). For other lineages, however, fluorescent protein expression varied depending on the mouse model, lineage, and even maturation step examined (Fig. 2A): for example, GFP expression was progressively lost during dendritic cell production and in specific dendritic cell subsets (up to 12% of plasmacytoid dendritic cells are GFP^-^), as well as eosinophils (Fig. 2B). In contrast all bone marrow dendritic progenitors, dendritic cell subsets and eosinophils were almost 100% TdTomato^+^. GFP expression was maintained throughout erythropoiesis while TdTomato expression was progressively lost during differentiation. In B cell differentiation up to 22% of common lymphoid progenitors (CLP) are TdTomato^-^ while some Pre-pro B, Pro B and Pre B cells lose GFP expression (Fig. 2B). Both strains lost around 10% fluorescent protein expression in basophils (Fig. 2B). Taking together these experiments demonstrate that expression of fluorescent reporter proteins in commonly used models varies in a cell-specific manner during hematopoiesis.

**Figure 2.**
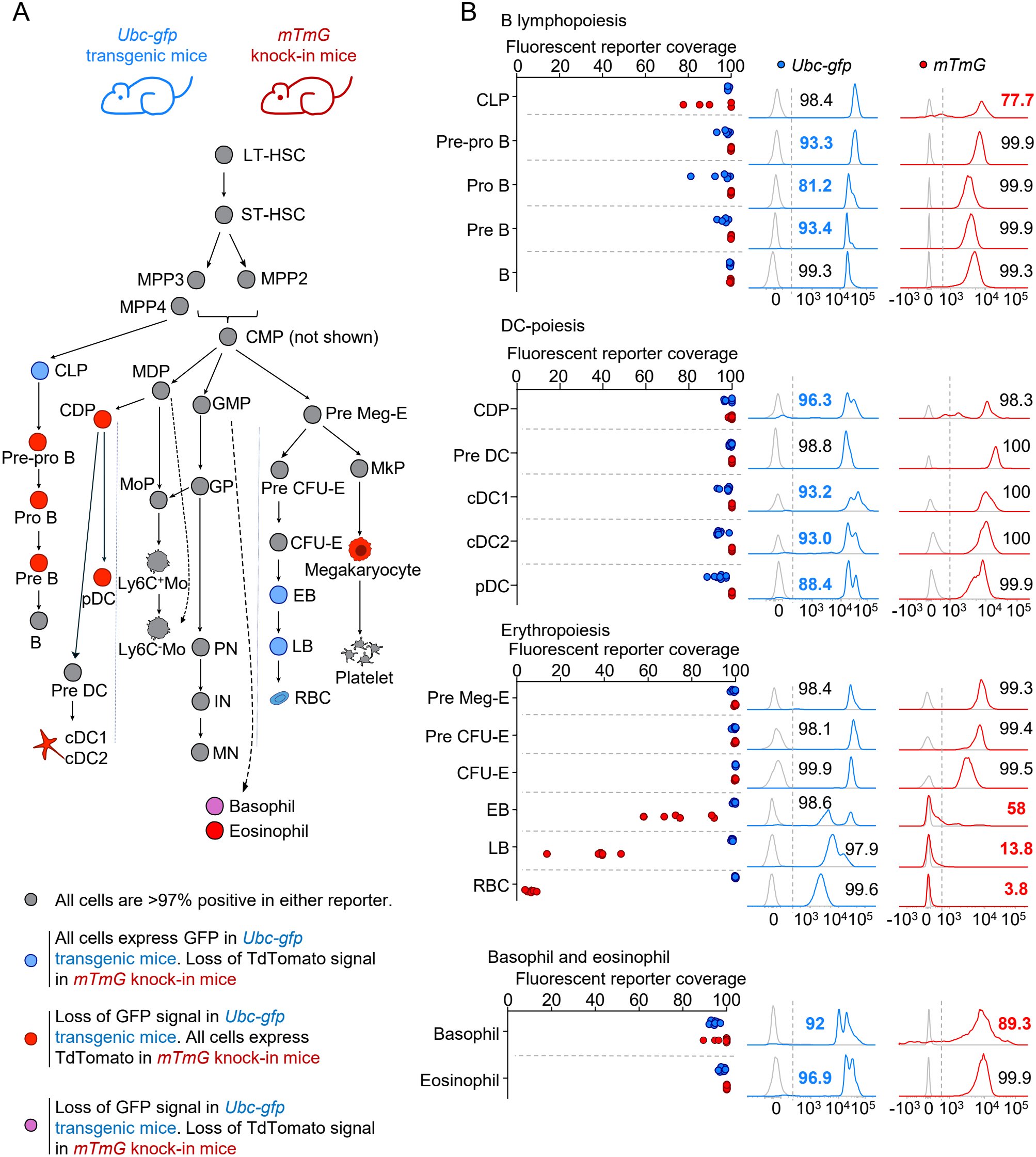
Heterogenous GFP and TdTomato expression at different stages of hematopoiesis in the bone marrow of *Ubc-gfp* and *mT/mG* mice. (A)Schematic representation of heterogeneous fluorescent protein expression at different stages of hematopoiesis. LT-HSC, long-term hematopoietic stem cell; ST-HSC, short-term hematopoietic stem cell; MPP, multipotent progenitor; CLP, common lymphoid progenitor; GMP, granulocyte-monocyte progenitor; GP, granulocyte progenitor; PN, promyelocyte; IN, immature neutrophil; MN, mature neutrophil; MDP, monocyte-dendritic cell progenitor; MoP, monocyte progenitor; Mo, monocyte; CDP, common dendritic cell progenitor; pDC, plasmacytoid dendritic cell; cDC, conventional dendritic cell; Pre-MegE, pre-megakaryocyte-erythroid progenitor; Pre-CFU-E, pre-colony-forming unit-erythroid; EB, erythroblast; LB, late erythroblast; Reti, reticulocyte; RBC, red blood cell; MkP, megakaryocyte progenitor. (B) Fluorescent expression (left panel) and MFI (right panel) in the indicated cells. In all panels expression of fluorescent proteins was detected by flow cytometry. Each dot represents one mouse in at least two independent experiments.

## Discussion

Fate mapping after adoptive transfer experiments requires precise tracking of the donor cells and their offspring *in vivo*. Here we have found that in *Ubc-gfp* mice, a mouse model widely used to track hematopoietic and immune cells after transfer, GFP is heterogeneously expressed in bone marrow hematopoietic cells, precluding its use for fate mapping of megakaryocytes, plasmacytoid dendritic cells and key steps of B lymphopoiesis. We have shown that these cell types can be precisely fate mapped via using *mTmG* mice as source of Tomato^+^ donor cells. However, this model shows heterogenous Tomato expression in common lymphoid progenitors and is progressively downregulated in erythroid cells. These findings highlight the importance of validating reporter protein expression in the cells of interest and provide a resource for fate mapping experiments of mouse bone marrow hematopoietic cells.

## Acknowledgments

We thank the Bioanalysis Imaging Facility, the Research Flow Cytometry Facility in the Division of Rheumatology, Veterinary Services, and the Cincinnati Children’s Single Cell Genomics Facility at Cincinnati Children’s Medical Center for experimental and technical assistance. This work was performed in Daniel Lucas lab in the United States and funded by a grant from the National Heart Lung and Blood Institute (R35HL177125 to DL.). DL is a Scholar of Blood Cancer United. Fees for research flow cytometry and imaging cores were partially subsidized by U54DK126108. Data generated using an BD/FACSymphony S6 cell sorter was funded by NIH grant S10OD023410.

## Figure Legend

**Supplementary Figure 1. Gating strategies for detection of hematopoietic cells by FACS**.

LT-HSC, long-term hematopoietic stem cell; ST-HSC, short-term hematopoietic stem cell; MPP, multipotent progenitor; CLP, common lymphoid progenitor; GMP, granulocyte-monocyte progenitor; GP, granulocyte progenitor; PN, promyelocyte; IN, immature neutrophil; MN, mature neutrophil; MDP, monocyte-dendritic cell progenitor; MoP, monocyte progenitor; Mo, monocyte; CDP, common dendritic cell progenitor; pDC, plasmacytoid dendritic cell; cDC, conventional dendritic cell; Pre-MegE, pre-megakaryocyte-erythroid progenitor; Pre-CFU-E, pre-colony-forming unit-erythroid; EB, erythroblast; LB, late erythroblast; Reti, reticulocyte; RBC, red blood cell; MkP, megakaryocyte progenitor.

**Supplementary Figure 2. Expression of GFP or TdTomato in HSC/MPP, granulo- and monopoiesis**.

(A-C) Fluorescent protein expression (left panel) and MFI (right panel) in BM HSC/MPP (A), granulo-(B) and monopoiesis (C). LT-HSC, long-term hematopoietic stem cell; ST-HSC, short-term hematopoietic stem cell; MPP, multipotent progenitor; GMP, granulocyte-monocyte progenitor; GP, granulocyte progenitor; PN, promyelocyte; IN, immature neutrophil; MN, mature neutrophil; MDP, monocyte-dendritic cell progenitor; MoP, monocyte progenitor; Mo, monocyte.

