## Supplementary Figures for "Variable expression of fluorescent protein expression in hematopoietic cells purified from ubiquitous reporter mice"

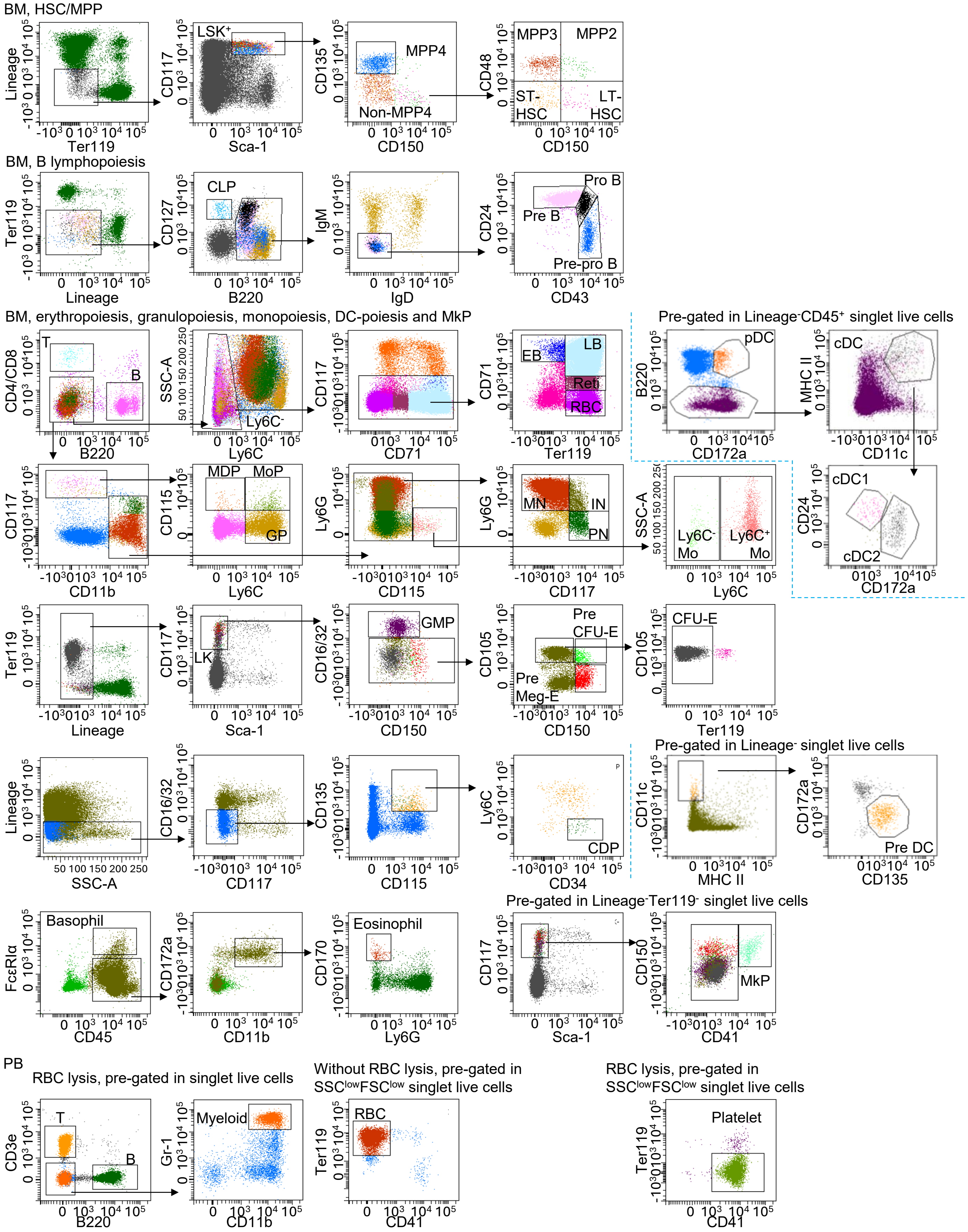


**Supplementary Figure 1. Gating strategies for detection of hematopoietic cells by FACS.**

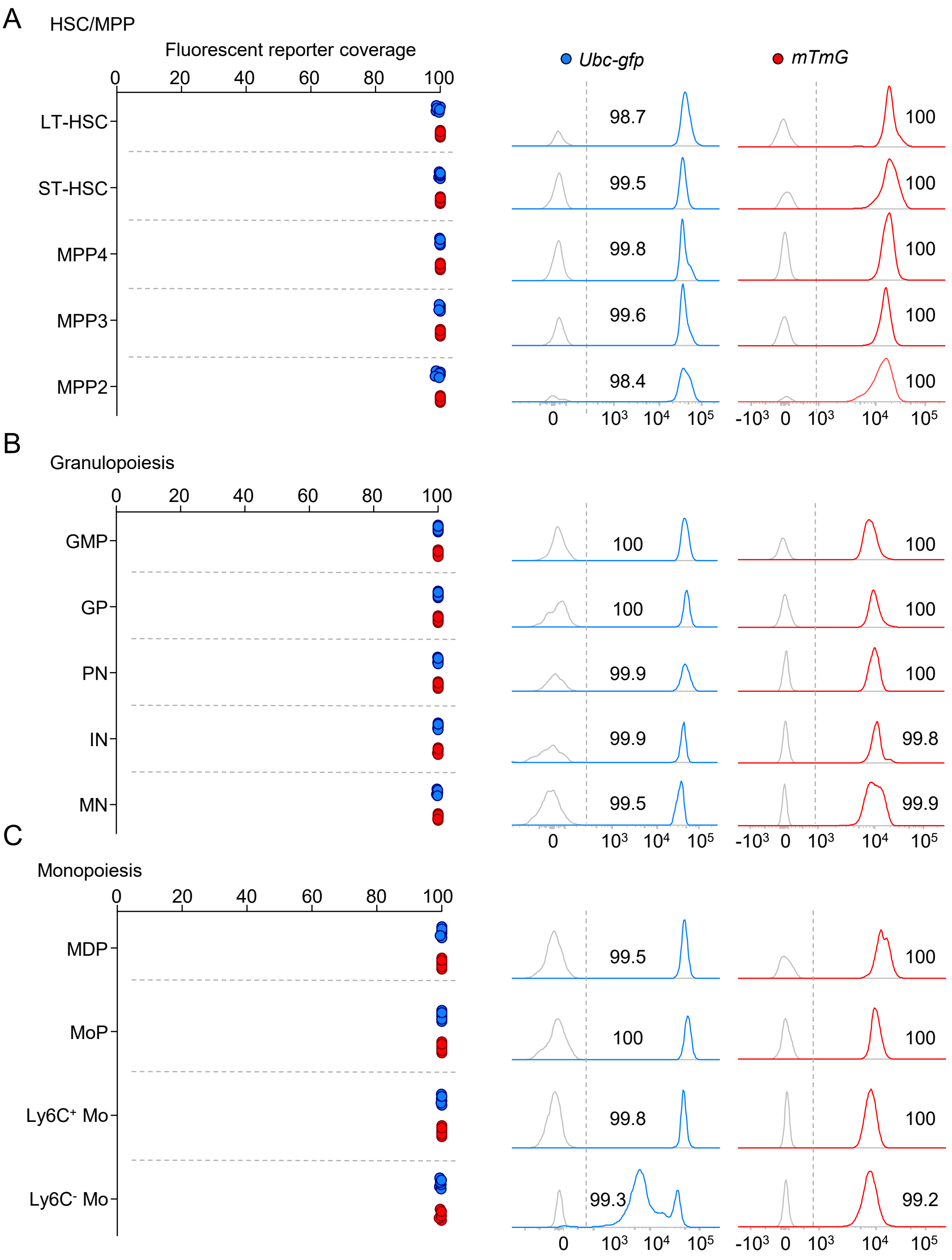


**Supplementary Figure 2. Expression of GFP or TdTomato in HSC/MPP, granulo- and monopoiesis.**
